# S100A4+ stromal cells are required for functional nipple development and successful nursing

**DOI:** 10.64898/2026.01.26.701676

**Authors:** Denisa Jaros Belisova, Ema Grofova, Viacheslav Zemlianski, Zuzana Sumbalova Koledova

## Abstract

The nipple and mammary gland are essential for the survival of mammalian offspring, providing postnatal nourishment. Their development, like that of epidermal appendages, depends on instructive signals from mesenchymal cells. S100A4 (S100 calcium binding protein A4, also known as fibroblast-specific protein 1) is expressed by mesenchymal cells and has been associated with hair follicle regeneration. S100A4-expressing cells have been implicated in the development of eccrine glands, and studies using *S100a4-Cre* to manipulate gene function have suggested that S100A4-expressing cells may contribute to mammary branching morphogenesis. However, the identity and functional contribution of S100A4-positive (S100A4+) cells to nipple and mammary development remain unclear. Here, we used a cell-depletion mouse model, *S100a4-Cre;DTA*, to investigate their role during lactation. *S100a4-Cre;DTA* dams exhibited a severe nursing defect leading to complete litter loss within the first day postpartum. Immunofluorescence analysis and oxytocin stimulation assays revealed no abnormalities in mammary morphology, milk production, or alveolar contractility, but defective nipple development was observed. Bulk RNA sequencing of nipple tissue indicated inflammatory signatures. Lineage tracing and immunofluorescence staining identified S100A4+ cells as fibroblasts and immune cells in the nipple, while in the mammary gland S100A4 was expressed only in immune cells. Our study uncovers a previously unrecognized role of S100A4+ cells in nipple development, highlighting their importance for successful lactation and offering new insights relevant to breastfeeding medicine.

## Introduction

During embryonic development, the epidermis originates from the surface ectoderm as a single layer and subsequently transforms into complex, multilayered skin^1^. This transformation is guided by the underlying mesenchyme, which is responsible for developing and maintaining the characteristics of different body regions, including epidermal appendages and areas of specialized epidermis^2,3^. The specialized epidermis diversifies from the trunk epidermis in order to mediate interactions with the environment and fulfil specific functions. To this end, it develops distinct morphological and molecular features, including the presence of epidermal appendages, a characteristic stratification pattern, and the expression of unique differentiation markers^4^. Specialized epidermis covers e.g. the anal and urogenital regions, palms, soles and lips in humans, and the tail, footpads, and muzzle in mice.

The nipple is another example of specialized epidermis that is anatomically and functionally connected to the mammary gland, an epidermal appendage^4,5^. The early embryonic development of both structures depends on the mammary mesenchyme^6,7^, which instructs the mammary placode to form the mammary epithelium and the overlying epidermis to become the nipple sheath^4,8–10^. These structures are unique to mammals and play an essential role in providing nourishment to offspring. Hormonal stimuli during pregnancy and lactation direct mammary alveogenesis, milk production and nipple enlargement, which are crucial for milk excretion and delivery to offspring^11,12^.

S100A4 (S100 calcium binding protein A4), also known as fibroblast-specific protein 1 (FSP1) or metastasin 1 (MTS1), is a member of the S100A calcium-binding protein family. It has been extensively studied in relation to tumor progression, metastasis, and fibrosis^13–16^. Nevertheless, its role in normal development is also increasingly recognized. S100A4 expression has been reported in embryonic mesenchyme^17,18^. S100A4-positive (S100A4+) dermal cells have been implicated in the development of the eccrine glands^19^, and S100A4 expression has been observed during hair follicle regeneration^20^. Postnatally, *S100a4* expression has been observed in various skin cell types, including Langerhans cells, melanocytes, keratinocytes, infiltrating lymphocytes, fibroblasts and fibrohistiocytes^19,21–23^. These findings indicate that S100A4 marks a heterogeneous population of cells in the skin. However, it remains unclear whether S100A4+ cells contribute to the development and function of the nipple-mammary gland complex.

In this study, we employ *S100a4-Cre;DTA* and *S100a4-Cre;mTmG* mouse models to investigate the role of S100A4+ cells in nipple and mammary gland development and function. We report an inverted nipple-like phenotype in *S100a4-Cre;DTA* dams and describe the distribution, morphology, and heterogeneity of S100A4+ cells in the mammary gland and nipple.

## Results

### The whole-litter mortality of S100a4-Cre;DTA mice is due to an impaired nursing capacity

To investigate the role of S100A4+ stromal cells, we used a genetic ablation model *S100a4-Cre;DTA*, in which Cre-driven expression of diphtheria toxin fragment A (DTA) induces cell-autonomous apoptosis in S100A4-expressing cells^24,25^. When *S100a4-Cre;DTA* and control *DTA* females were bred with wild-type males, a slightly lower number of pups were born to *S100a4-Cre;DTA* dams (Figure 1a). Examination of the dams after parturition *post mortem* did not reveal any remaining pups *in utero*; therefore, the decreased number of pups was not caused by complications during birth. The dams displayed normal maternal behavior, including nesting. However, no milk intake was observed in the litters of *S100a4-Cre;DTA* dams, demonstrated as an absence of the milk spot in their pups (Figure 1b). The lack of nutrition in pups from *S100a4-Cre;DTA* dams resulted in early litter mortality on lactation day 1 (L1), with no pups surviving to lactation day 2 (L2). Fostering experiments revealed normal survival of the pups of *S100a4-Cre;DTA* dams when nursed by control mothers (Figure 1c), suggesting that the early litter mortality of pups from *S100a4-Cre;DTA* mothers is caused by a nursing defect in lactating dams rather than health issues in the pups themselves.

**Figure 1.**
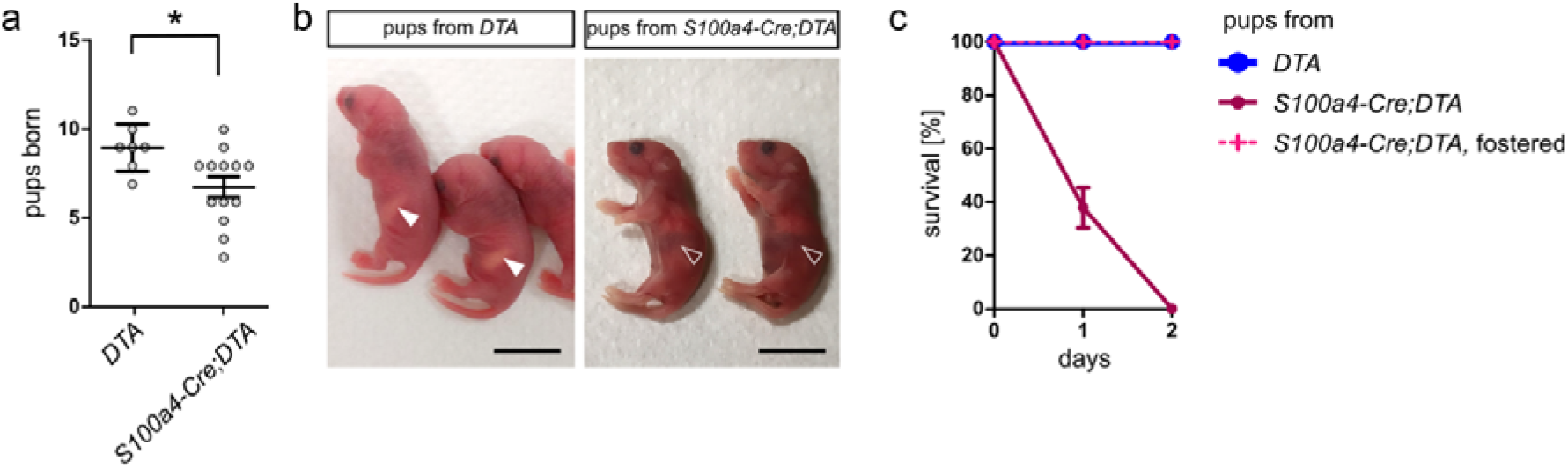
The whole-litter mortality of *S100a4-Cre;DTA* mice is a result of impaired nursing capacity. (a) Number of pups born to *DTA* and *S100a4-Cre;DTA* dams. The plot shows the mean ± SD, *p < 0.05 (Mann-Whitney test), n = 5 *DTA*/16 *S100a4-Cre;DTA.* (b) Pictures of the pups from *DTA* and *S100a4-Cre;DTA* dams. Arrowheads indicate milk spots. Scale bar = 1 cm. (c) Quantification of litter survival on L1 and L2. n = 3 *DTA*/3 *S100a4-Cre;DTA*/1 *S100a4-Cre;DTA* fostered.

### The ablation of S100A4+ cells delays the branching morphogenesis of the mammary gland during puberty, but this phenotype is resolved in adulthood

To investigate the potential contribution of systemic or local mammary developmental defects to the lactation defect, we compared body weight and mammary epithelial development between *S100a4-Cre;DTA* mice and *DTA* littermates. We observed significantly lower body weight in *S100a4-Cre;DTA* mice (Figure 2a), consistent with the previously reported function of S100A4+ fibroblasts in the adipogenic niche^26^. However, we found no difference in the size of the mammary fat pad nor in the size of adipocytes between the *S100a4-Cre;DTA* mice and *DTA* controls (Figure 2b,c, S1a,b). Immunofluorescent examination of epithelial organization showed no detectable abnormalities in *S100a4-Cre;DTA* mammary glands (Figure 2c).

**Figure 2.**
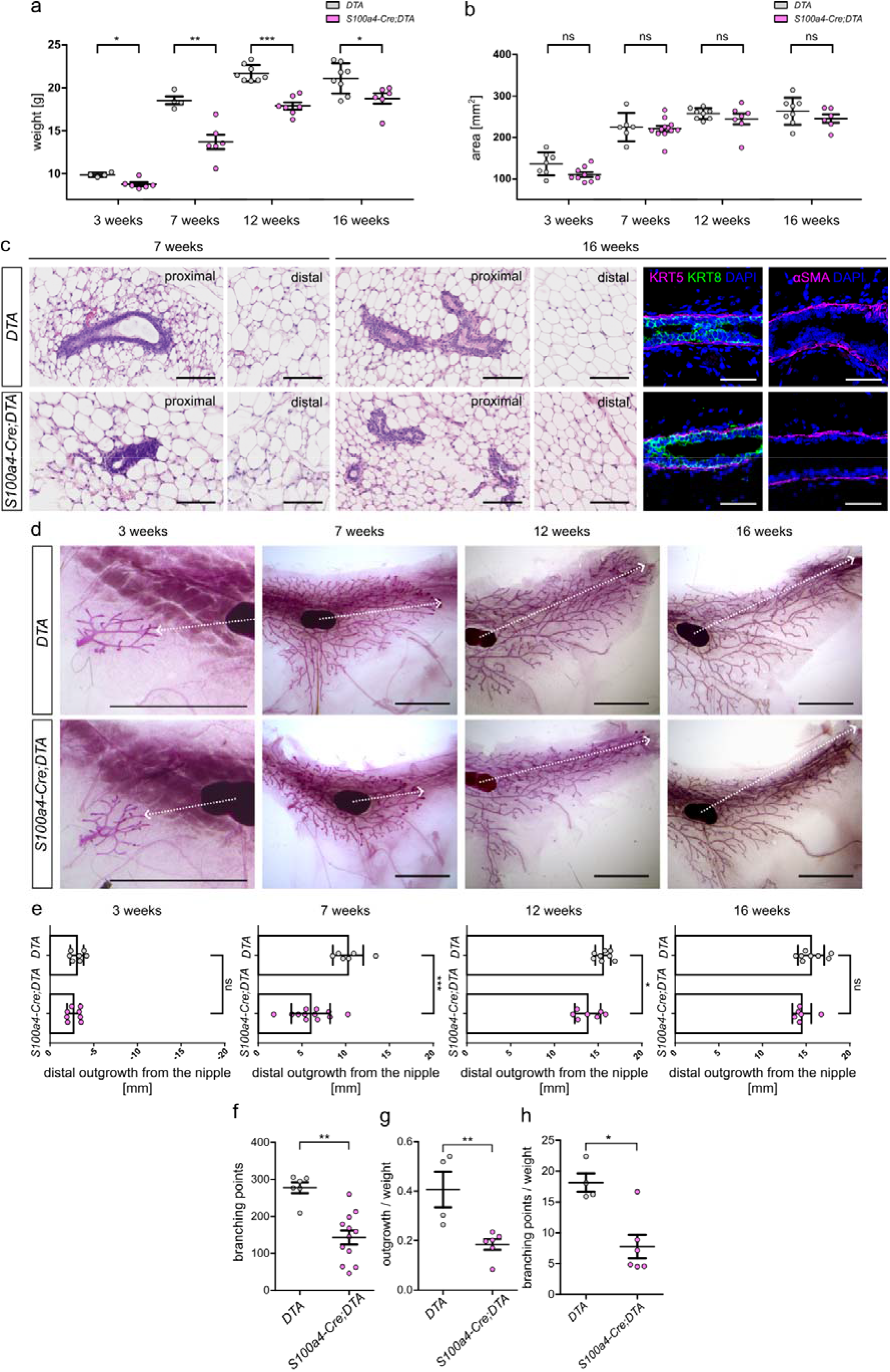
Ablation of S100A4+ cells delays pubertal branching morphogenesis and the phenotype is resolved in adulthood. (a, b) Quantification of (a) body weight, measured as the weight of the mouse *post mortem*, and (b) mammary fat pad size, measured as the area of the fat pad on carmine-stained whole-mount mammary glands, in *DTA* and *S100a4-Cre;DTA* mice at 3, 7, 12 and 14 weeks of age. The plots show the mean ± SD, ns p > 0.05, *p < 0.05, **p < 0.01, ***p < 0.001 (Mann-Whitney test), n = 4/3/2/3 litters/time-point, N = 7/12; 4/6 (a) or 6/12 (b); 8/7; 8/6 for *DTA/S100a4-Cre;DTA* mice and respective time-points. (c) Hematoxylin and eosin-stained paraffin sections of the mammary fat pad at 7 and 16 weeks of age and representative pictures of immunostaining for KRT5 (myoepithelial cells) and KRT8 (luminal cells); and αSMA in *DTA* and *S100a4-Cre;DTA* mammary glands at 16 weeks of age. Scale bar (c,d) = 100 µm. (d) Representative phenotypes of *DTA* and *S100a4-Cre;DTA* carmine-stained whole-mount mammary glands. Scale bar = 5 mm. (e) Quantification of epithelial outgrowths distally from the lymph node. (f-h) The plots show total number of branching points (f), epithelial outgrowth [mm] normalized to body weight [g] (g), and total number of the branching points normalized to body weight [g] (h) in 7 weeks old *DTA* and *S100a4-Cre;DTA* mice. All plots show the mean ± SD, ns p > 0.05, *p < 0.05, **p < 0.01, ***p < 0.001 (Mann-Whitney test), n = 4/3/2/3 litters/time-point, N = 7/12; 6/12 (e, f) or 4/6 (g, h); 8/7; 8/6 for *DTA/S100a4-Cre;DTA* mice and respective time-points.

Analysis of mammary epithelial development using whole-mount carmine staining revealed no significant differences in the prenatal establishment of the mammary epithelial tree, but it did reveal significantly delayed epithelial outgrowth and reduced branching in pubertal (7 weeks old) *S100a4-Cre;DTA* mice (Figure 2d-f). Normalization of epithelial outgrowth and branching to body weight indicates that the observed defect represents a mammary-specific impairment rather than a consequence of reduced body growth (Figure 2g,h). However, this phenotype was transient, with the epithelium in *S100a4-Cre;DTA* mice achieving the same length in the adulthood (16 weeks) as in the *DTA* mice (Figure 2d,e). Overall, *S100a4-Cre;DTA* mice are smaller and exhibit a transient delay in mammary epithelial morphogenesis during puberty; however, the epithelial outgrowth is restored in adulthood.

### The mammary glands of S100a4-Cre;DTA lactating dams exhibit normal histology, yet show signs of milk stasis

In order to investigate the mechanism of the nursing defect in *S100a4-Cre;DTA* lactating dams, we examined the architecture and histology of the lactating mammary glands of *S100a4-Cre;DTA* and *DTA* mice. Macroscopically, we observed a significant accumulation of milk in the lactiferous duct of the *S100a4-Cre;DTA* mammary glands (Figure 3a), which indicates milk stasis. Under normal conditions, such as in control *DTA* mice, the milk does not accumulate in the duct because it is regularly suckled by the pups. Further investigation of carmine-stained whole-mount mammary glands revealed no difference in alveolar formation between the *S100a4-Cre;DTA* and *DTA* mice (Figure 3b).

**Figure 3.**
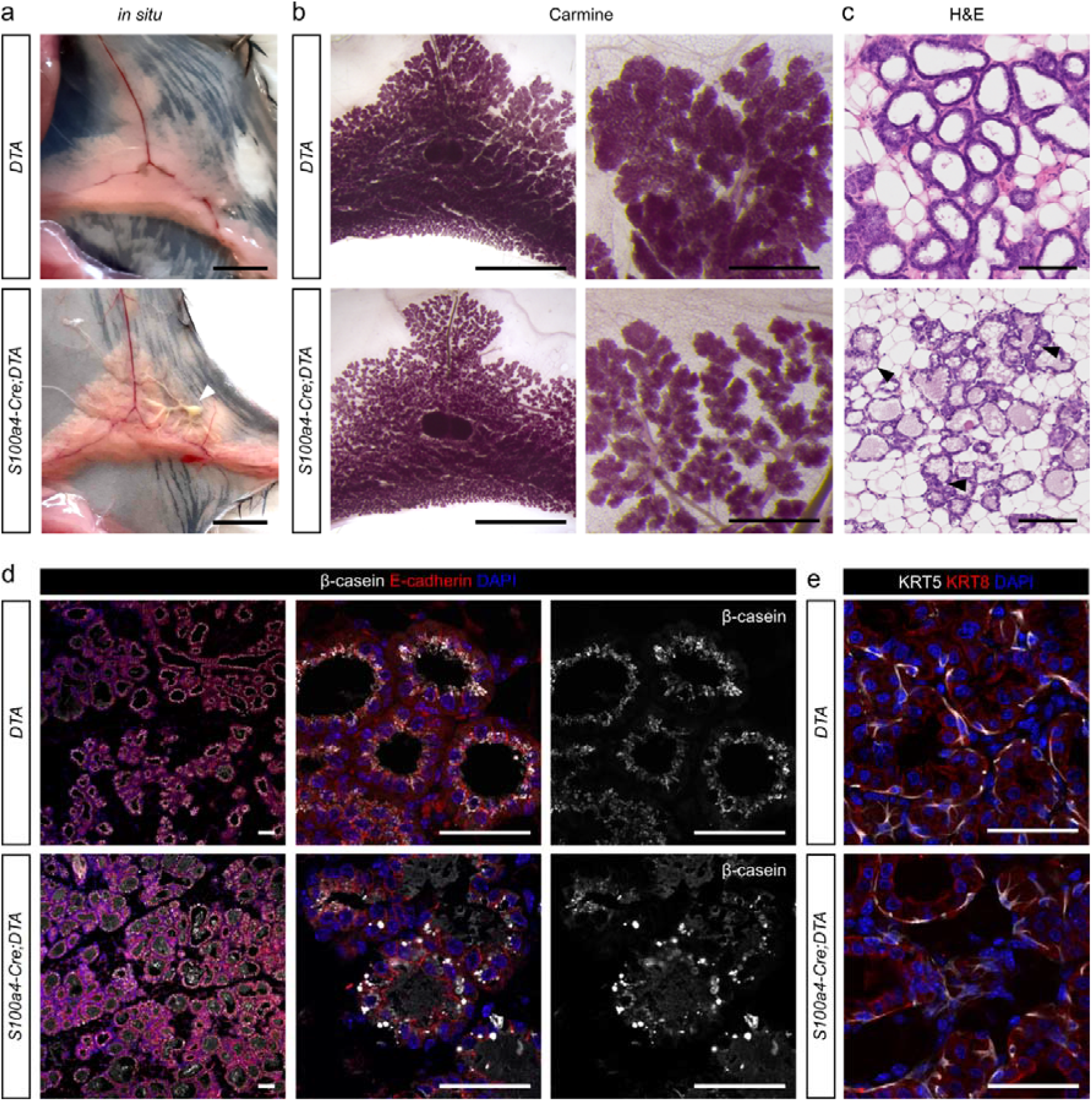
The mammary tissue of *S100a4-Cre;DTA* lactating dams exhibits normal histology but shows signs of milk stasis. (a) Macroscopic images of the inguinal mammary glands *in situ*. White arrowhead indicates lactiferous duct filled with milk. Scale bar = 5 mm. (b) Representative photographs of carmine-stained mammary glands from lactating (L1) *DTA* and *S100a4-Cre;DTA* mice. Scale bar = 5 mm and 1 mm (magnification). (c) H&E-stained paraffin sections of lactating (L1) mammary glands from *DTA* and *S100a4-Cre;DTA* mice. Black arrowheads indicate cytoplasmic lipid droplets. Scale bar = 100 µm. (d, e) Representative pictures of immunostaining for β-casein and E-cadherin (epithelium) (d) and KRT5 (myoepithelial cells) and KRT8 (luminal cells) (e). Scale bar = 50 µm.

However, analysis of tissue sections revealed the accumulation of a pink-stained substance (presumably milk) within the alveoli, as well as the presence of cytoplasmic lipid droplets, in *S100a4-Cre;DTA* mice (Figure 3c). This suggests a nursing defect^11,27^. To verify the presence of milk in the alveoli, we performed immunofluorescent staining for β-casein. In control *DTA* mammary glands, β-casein was localized apically in luminal cells and minimally in the alveolar lumen. However, in *S100a4-Cre;DTA* mammary glands, the apical localization of β-casein in luminal cells was disrupted, with β-casein accumulating in the alveolar lumen instead (Figure 3d). Immunolabeling for myoepithelial (KRT5) and luminal (KRT8) cells revealed physiological cellular architecture in both *S100a4-Cre;DTA* and *DTA* mammary glands (Figure 3e).

Lactation is tightly controlled by two key hormones, prolactin and oxytocin, which are produced or stored in the pituitary gland. A previous study reported on the importance of S100A4+ pituitary hematopoietic cells in regulating fertility and endocrine function^28^. Therefore, we investigated whether the pituitary gland defect could contribute to the impaired nursing capacity of *S100a4-Cre;DTA* dams. We observed S100A4-lineage-traced cells in the pituitary gland and ovaries of *S100a4-Cre;mT/mG* mice (Figure S2a,b). However, we found no macroscopic or microscopic aberrations in *S100a4-Cre;DTA* pituitary glands (Figure S2c,d). Moreover, examination of the estrous cycle and ovaries revealed no significant differences between *S100a4-Cre;DTA* and *DTA* females (Figure S2e,f), suggesting that a systemic hormonal disbalance is unlikely to be the underlying cause of the observed phenotype in *S100a4-Cre;DTA* dams.

To test the milk-ejection function of the alveoli, including their responsiveness to oxytocin and contractile capacity, we performed an *ex vivo* oxytocin contraction assay. In this assay, we exposed lactating mammary tissue (L1) to oxytocin stimulation under time-lapse imaging surveillance. Both *DTA* and *S100a4-Cre;DTA* alveoli contracted in response to oxytocin (Figure S3, Movies S1 and S2), indicating that there was no defect in the response to exogenous oxytocin. Together, these data demonstrate that the *S100a4-Cre;DTA* mice form functional lactating alveoli, but display signs of milk stasis, indicating impaired nursing capacity in *S100a4-Cre;DTA* dams.

### The depletion of S100A4+ stromal cells results in abnormal nipple development

The ability to nurse relies on the lactation capacity of the mammary glands to produce adequate milk, as well as the functionality of the nipple to deliver milk effectively to offspring. As there was no evidence of impaired lactation capacity, we investigated nipples in the *S100a4-Cre;DTA* and *DTA* mice. In *DTA* females, the nipple protruded normally and became more prominent during lactation (Figure 4a). However, the nipple in *S100a4-Cre;DTA* females was significantly smaller and did not protrude or achieve the typical conical shape (Figure 4a, Figure S4c). Longitudinal sections through the nipples revealed a major developmental defect in *S100a4-Cre;DTA* nipples, including a lack of protrusive growth and dermal sheath coverage (Figure 4b, c). The *S100a4-Cre;DTA* nipple phenotype somewhat resembled an inverted nipple, which can be manually exposed from its depressed position. Similarly, applying pressure to the surrounding skin exposed the abnormal nipple in *S100a4-Cre;DTA* mice (Figure 4d).

**Figure 4.**
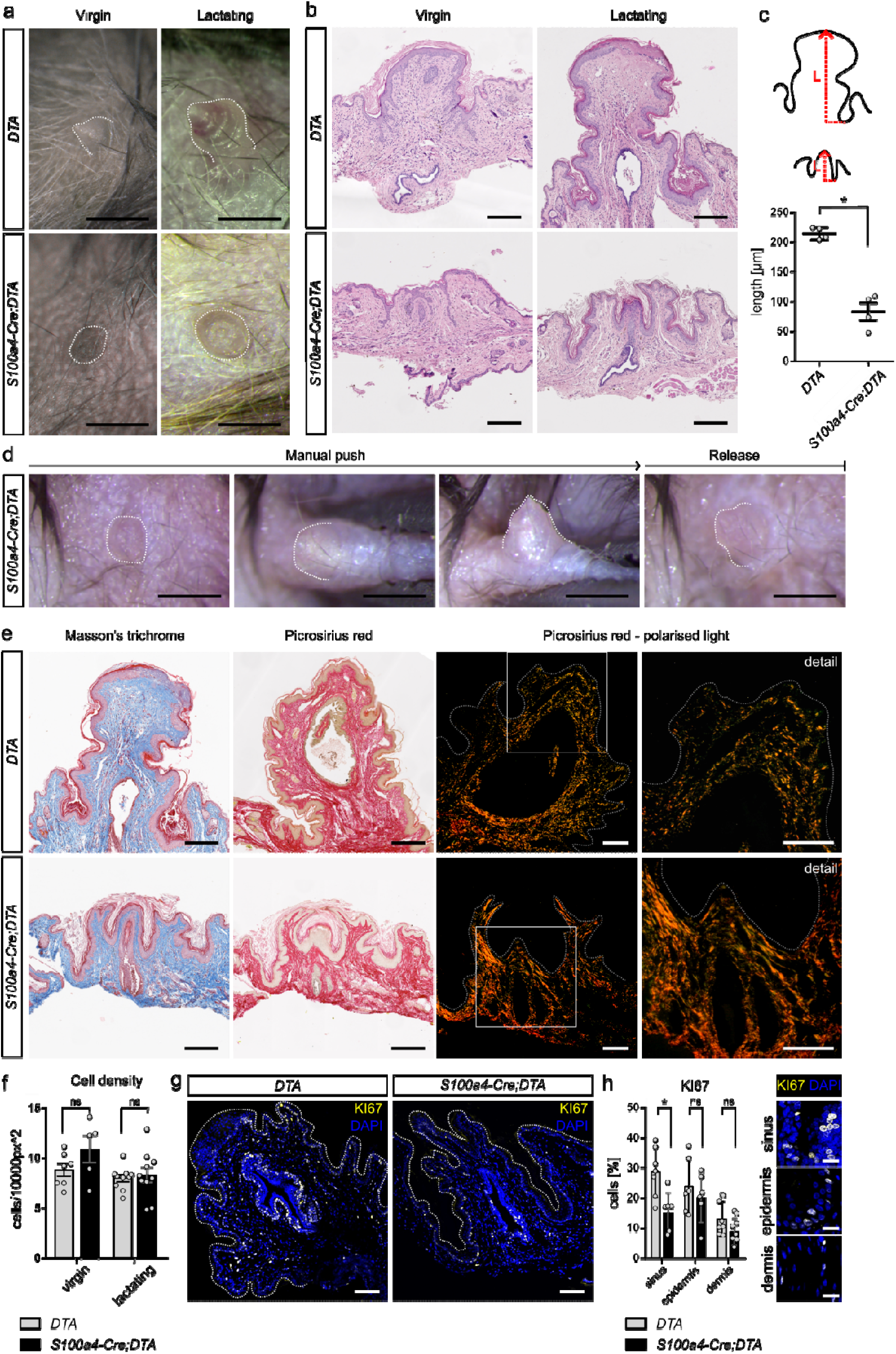
Depletion of S100A4+ stromal cells results in abnormal nipple development. (a) Representative photographs of nipples from *DTA* and *S100a4-Cre;DTA* virgin (12+ weeks old) and lactating (L1) dams *in situ.* Scale bar = 1 mm. (b) H&E-stained tissue sections of the nipples. Scale bar = 200 µm. (c) Quantification of nipple length (L), which is measured as the length of the nipple from the nipple-areola ridge to the most distant nipple point and respective scheme of the measurement. The plot shows the mean ± SD, *P < 0.05 (Mann-Whitney test), n = 4 *DTA* / 4 *S100a4-Cre;DTA*. (d) Photodocumented process of the inverted nipple exposure, and end-point after manual push release. Scale bar = 1 mm. (e) Representative pictures of histological sections of *DTA* and *S100a4-Cre;DTA* nipples stained for collagen by Masson’s trichrome or Picrosirius red. Scale bar = 200 µm. The right panel shows visualization of collagen organization and structure by polarized light microscopy of Picrosirius red-stained sections. Typically, thicker collagen fibers exhibit stronger birefringence and appear red or orange, while thinner fibers exhibit weaker birefringence and appear green or yellow. The white boxes mark the area depicted in detail picture. Scale bar = 100 µm. (f) Quantification of cell density in the dermis of virgin and lactating (L1) nipples, measured as the number of nuclei per 10,000 px^2^. The plot shows the mean ± SD, ns p > 0.05, n = 4 *DTA* / 3 *S100a4-Cre;DTA* virgin, 4 *DTA* / 5 *S100a4-Cre;DTA* lactating, N = 5 – 10 FOV for both genotypes and time points. (g) Immunofluorescent labeling for KI67 in lactating (L1) nipples. Scale bar = 100 µm. In (a, d, e, g) dotted lines indicate nipple border. (h) Quantification of cell proliferation in various nipple areas (L1): lactiferous sinus, epidermis and dermis as shown in the legend, scale bar = 20 µm; measured as percentage of KI67+ cells to the total number of cells in the area. The plot shows the mean ± SD, ns p > 0.05, *p < 0.05, n = 3 *DTA* / 3 *S100a4-Cre;DTA,* N = 6 – 8 FOV per nipple area for both genotypes.

Morphological changes to the nipple induced by pregnancy and lactation include epidermal cell proliferation and changes to the connective tissue, particularly increased collagen and elastic fiber deposition^12,29^. Analysis of collagen composition and organization (using Masson’s trichrome and picrosirius red staining) suggested that the collagen fibers were loosened in the *DTA* nipples and densely packed in the *S100a4-Cre;DTA* nipples (Figure 4e, Figure S5a). However, the quantification (mean intensity value for mature collagen and CT-FIRE analysis) failed to reveal significant difference between controls and mutants (Figure S5b-e). Proliferation analysis revealed decreased proliferation in the *S100a4-Cre;DTA* L1 nipple, with significantly less proliferation in the lactiferous sinus of the *S100a4-Cre;DTA* L1 nipple than in the *DTA* L1 nipple (Figure 4g,h). There were no abnormalities in epithelial organization, smooth muscle along the lactiferous duct, or sebaceous glands in *the S100a4-Cre;DTA* L1 nipple (Figure S6a-c). Taken together, these data suggest that the observed nursing failure in *S100a4-Cre;DTA* dams is caused by abnormal nipples that lack protrusion and cannot therefore be grasped and suckled by the pups. Abnormal nipple development in *S100a4-Cre;DTA* dams is associated with abnormal collagen deposition and insufficient dermal proliferation.

### S100A4+ cells comprise fibroblasts and immune cells

To investigate the morphology and distribution of S100A4+ cells and their progeny, we used the *S100a4-Cre;mT/mG* reporter mouse model. In this model all cells express membrane Tomato (mT; a red fluorescent protein) unless they express Cre under the control of the *S100a4* promoter. This switches the expression from membrane Tomato to membrane green fluorescent protein (mG; GFP)^30^. This genetic switch in cells is permanent and is passed on to the cell progeny, enabling lineage tracing.

Using the optical tissue clearing technique CUBIC^31^ combined with confocal imaging, we visualized *S100a4*-labelled GFP+ cells in the nipple at different developmental stages. In embryonic skin tissue (E15.5 and E18.5), we observed a small number of rounded GFP+ cells in the nipple sheath (Figure 5a, Figure S4a). In the adult virgin nipple (10 weeks old), both rounded and spindle-shaped GFP+ cells were localized concentrically around the lactiferous duct and formed a significant proportion of the nipple dermis. During lactation, when the nipple increases in size, the GFP+ stromal component becomes even more prevalent (Figure 5a). In the mammary gland, however, the majority of GFP+ cells had a rounded morphology. In embryonic tissue, they were almost exclusively dispersed in the mesenchyme. In the adult virgin or lactating tissue, the GFP+ cells were spread throughout the fat pad but were also localized in the periepithelial stroma and infiltrated the epithelium (Figure S7a,b). We also observed GFP+ cells in close association with the endothelium, consistent with an immune cell lineage (Figure 5b,c). Immunofluorescent analysis revealed GFP+ S100A4+ cells surrounding the mammary epithelium (Figure S7c). In contrast, GFP+ cells co-expressing the fibroblast marker PDGFRα were rare or nearly absent (Figure S7d), whereas GFP+ F4/80+ cells were frequently observed (Figure S7e). In the skin, *S100a4* is expressed in immune cells and fibroblasts^32^. Analysis of cells acutely expressing S100A4, as well as cells derived from S100A4-expressing cells, by co-labeling GFP and S100A4, revealed that S100A4-GFP+ cells contributed to the epidermal cell lineage. In contrast, S100A4+ GFP+ cells were restricted to dermis or were intercalated between epidermal cells. These cells can be identified as Langerhans cells due to their morphology and expression of AIF1 (Figure 5d,f). To investigate the nature of the GFP+ cells in the nipples of *S100a4-Cre;mT/mG* females, we performed an immunofluorescence analysis, co-staining for GFP and immune cell markers (AIF1, CD45) and the fibroblasts markers (VIM, PDGFRα) and performed the respective quantification (Figure 5e). We found that GFP+ cells co-expressing immune markers had a rounded morphology with protrusions and often penetrated the epidermis or the epithelium of the lactiferous ducts (Figure 5f,g) whereas GFP+ cells co-expressing fibroblast markers were localized in the dermis and had spindle-shaped morphology and are more prevalent (Figure 5e,h,i). Overall, our data suggest that S100A4-expressing cells in the nipple comprise immune cells and fibroblasts. Interestingly, S100A4 antibody labeling revealed presence of S100A4+ cells in *S100a4-Cre;DTA* tissues (Figure S4b, Figure S8a,b).

**Figure 5.**
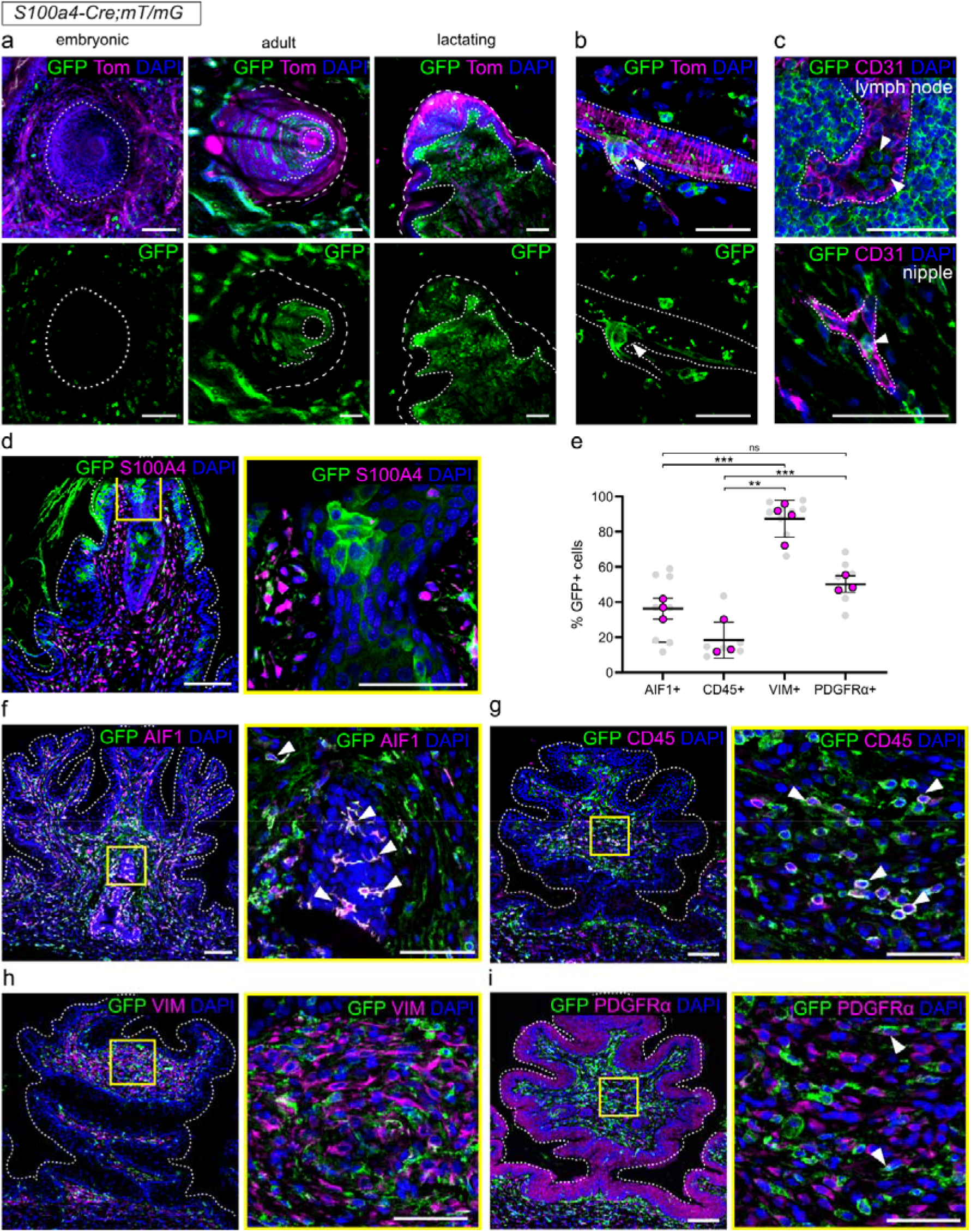
S100A4+ lineage cells are heterogeneous, comprising fibroblasts and immune cells. (a) Representative images of cleared whole-mount *S100a4-Cre;mT/mG* nipple tissue at different developmental time-points: embryonic (E18.5), adult virgin (10 weeks of age), lactating (L1). Dashed line delineates the nipple, dotted line delineates the nipple stroma. (b) A cryosection of *S100a4-Cre;mT/mG* tissue, showing GFP+ cell (white arrowhead) that has penetrated the endothelium (dotted line). (c) Immunofluorescent co-labeling of GFP and CD31 in the lymph node and nipple stroma showing GFP+ lineage cells (white arrowheads) inside the vasculature (dotted line). (d) Immunofluorescent co-labeling of GFP and S100A4 in the *S100a4-Cre;mT/mG* nipple. (e) Quantification of the contribution of GFP cells to immune cell or fibroblast lineages, calculated as the percentage of double-positive cells among all GFP cells. The plot shows the mean ± SD; Statistical significance was assessed using one-was ANOVA followed by Tukey’s multiple-comparison test; ns p > 0.05, **p < 0.01, ***p < 0.001, n = 3 – 4 *S100a4-Cre;mT/mG* animals (pink dots), N = 2 - 4 tissue section per animal (grey dots). (f - i) Immunofluorescent co-labeling of GFP with immune cell markers, AIF4 (f) and CD45 (g), and fibroblasts markers, vimentin (h) and PDGFRα (i), on FFPE sections of *S100a4-Cre;mT/mG* nipple tissue. Dotted lines outline the nipple tissue, yellow boxes mark the area depicted in the detailed picture of higher magnification. Scale bar = 50 µm (details in yellow boxes) and 100 µm.

### Transcriptome of S100a4-Cre;DTA nipple indicates concurrent tissue inflammation and regeneration

To investigate differences in transcription between *S100a4-Cre;DTA* and *DTA* nipples, we performed bulk RNA sequencing on nipples isolated from lactating dams (L1). This revealed 474 differentially expressed genes, of which 293 were significantly upregulated and 181 were significantly downregulated in *S100a4-Cre;DTA* nipples. The upregulated genes were associated with the metabolism of xenobiotics, including bacterial toxins (*Pon1* and various cytochromes, *Cyp2a5, Cyp2g1, Cyp2j7*), inflammation and immune cell recruitment (*Ccl20, Cxcl10, Il20ra, Tnfsf11, Serpina3b, Serpina3j*), the lipid skin barrier (*Elovl3, Elovl5, Elovl6*) and skin regeneration (*Krt17, Krt75, Egr1, Fbn2, Adam23*). The downregulated genes suggested a decrease in the luminal differentiation/secretory mammary program (*Wap, Csn1s1, Krt8, Krt18, Krt19, Muc4, Aqp5*) and altered ECM remodeling (*Lyve1, Tnc, Adamts5, Col7a1, Ecm1, Timp3*) (Figure 6a). This was consistent with the altered ECM organization observed in *S100a4-Cre;DTA* nipples. Gene ontology (GO) functional enrichment analysis clustered upregulated genes into the groups and identified the biological processes that were upregulated in *S100a4-Cre;DTA* nipples. The GO terms indicated that the tissue reacted to a foreign chemical or an endogenous compound (xenobiotic metabolic process, cellular response to xenobiotic stimulus, response to xenobiotic stimulus, epoxygenase P450 pathway), and responded to inflammation and repair (actin filament-based process, actin cytoskeleton organization; eicosanoid and lipid metabolic processes) (Figure 6b).The obtained data suggest that the *S100a4-Cre;DTA* nipple tissue responds to DTA expression and regenerates in order to cope with the loss of depleted cells.

**Figure 6.**
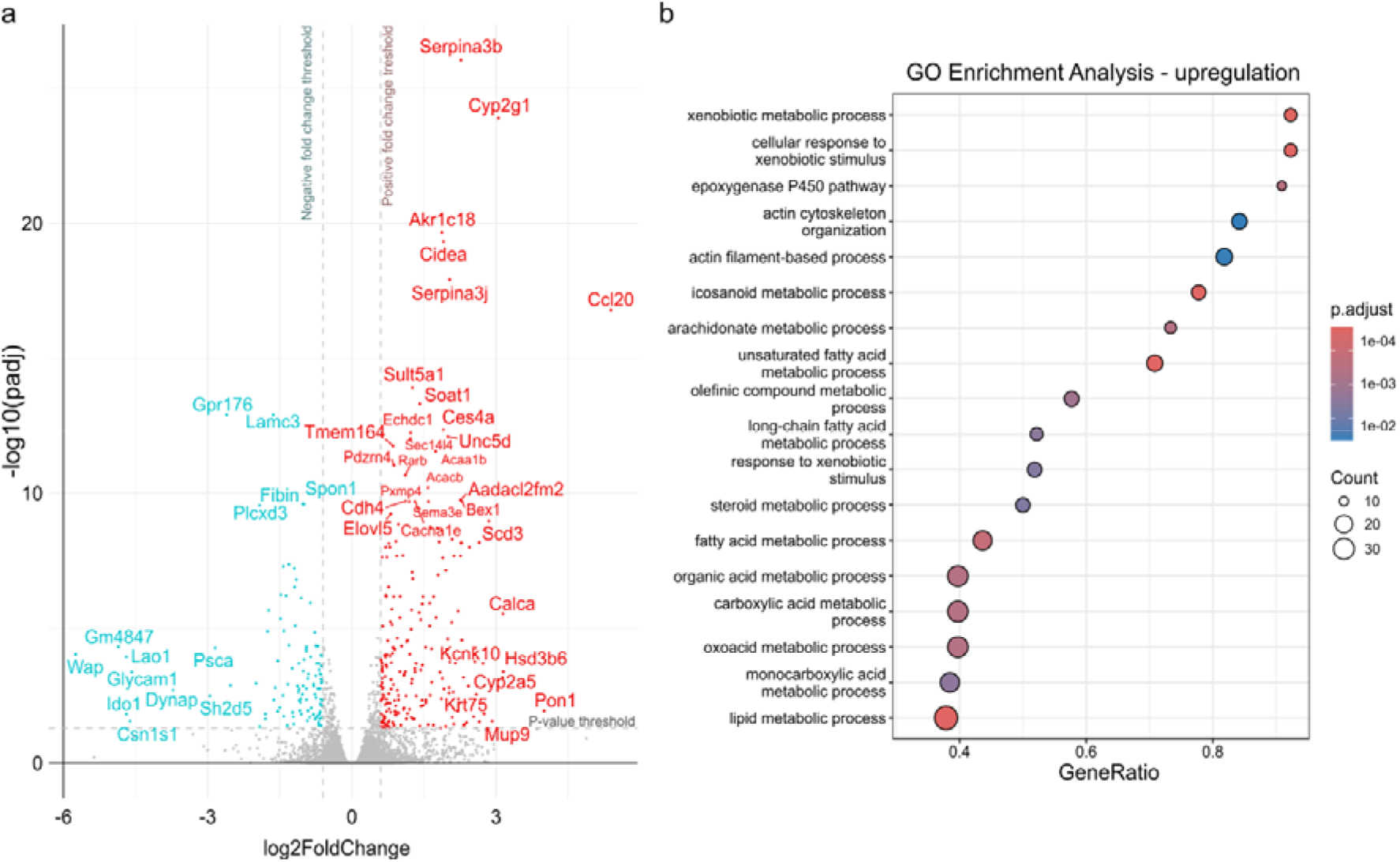
The transcriptome of *S100a4-Cre;DTA* nipples indicates concurrent tissue inflammation and regeneration. (a) A volcano plot visualizing the identified differentially expressed genes (DEGs). (b) Gene ontology enrichment analysis of the upregulated biological processes.

## Discussion

In this study, we identify a previously unrecognized role of S100A4+ stromal cells in nipple development using the *S100a4-Cre;DTA* mouse model. We demonstrate that depletion of S100A4-expressing cells results in nipple deformity that leads to nursing failure. Our data further indicate that S100A4+ cells represent a heterogeneous stromal population comprising both fibroblasts and immune cells, consistent with growing evidence that S100A4 expression is not restricted to a single lineage.

The observed lactation defect initially suggested impaired mammary gland function. However, systemic analysis of mammary development revealed only a transient delay in branching morphogenesis during puberty. Importantly, this phenotype resolved in adulthood, with normal epithelial development and the ability to undergo alveologenesis during pregnancy. These findings indicate that the primary cause of nursing defect in this model is unlikely to be a persistent defect in the mammary epithelium.

Instead, we identify abnormal nipple development as the principal defect. Nipple formation is initiated during embryogenesis under the control of the mammary mesenchyme^8,33^, and postnatal remodeling of the nipple depends on epithelial-stromal interactions as well as hormonal and mechanical cues^12,29^. In *S100a4-Cre;DTA* females, nipples fail to protrude and remain covered by dermal sheets, resembling features of inverted nipple, a condition in which the nipple does not project beyond the surface of the areola^34^. Given that proper nipple morphology, particularly its enlargement, is essential for effective suckling^35,36^, this structural defect provides a direct explanation for the observed nursing failure and is consistent with the fact that inverted nipples can cause breastfeeding difficulties^37,38^.

Our findings are also consistent with earlier hypotheses proposing that congenital inverted nipple may arise from insufficient development or remodeling of the underlying mesenchyme, which is consequently unable to push the nipple out of its depressed position^39^. While clinical work has largely focused on surgical correction, the developmental and cellular mechanisms underlying this condition have remained poorly understood. Our data provide experimental evidence that stromal cell populations are critical regulators of nipple morphogenesis.

Stromal cells, including fibroblasts and immune cells such as macrophages, play a role in the morphogenesis and maintenance of skin derivates, such as mammary gland, and specialized skin, such as the nipple^13,40–42^. In our cell-specific depletion study, we describe S100A4+ cells in the nipple and identify them as a heterogeneous population of cell comprising fibroblasts and immune cells. However, we did not observe fibroblast-like cells in the mammary gland. This aligns with growing evidence disputing the specificity of the S100A4 marker. While originally proposed as a great marker of fibroblasts in tissue^18,43^, it is being recognized that S100A4 is expressed by other cell types, including myeloid cells, rather than fibroblasts^14,28,44,45^. There is a prominent population of S100A4+ cancer-associated fibroblasts in breast cancer, while no S100A4+ fibroblasts are present in the normal mammary fat pad^16^. There, *S100A4* expression was specifically found in cells of hematopoietic origin^46^.In healthy skin, *S100A4* expression was identified in macrophages, T cells and dendritic cells and fibroblasts^32^, what aligns with our observation. S100A4 is also expressed in the developing eccrine gland and hair follicle^19–21^.

A key consideration when interpreting studies involving S100A4 is that fundamentally different experimental approaches have been used to investigate its role. These include descriptive analyses of S100A4 expression, functional studies targeting the S100A4 protein itself, genetic models using *S100a4-Cre* to manipulate unrelated genes in S100A4-expressing cells, and ablation models such as *S100a4-Cre;DTA*, which deplete S100A4 cells. These approaches are not equivalent and provide distinct types of information. In the present study, we specifically assess the consequences of ablating S100A4-expressing cells, and comparisons to other studies should therefore be interpreted within this context.

Studies using *S100a4-Cre* to manipulate specific signaling pathways (e.g. Wnt or Hedgehog signaling via gene deletion) in S100A4-expressing cells have reported diverse phenotypes, including effects on fertility and endocrine function^28,44^. However, these phenotypes primarily reflect the consequences of pathway perturbations within S100A4-expressing cells rather than the role of S100A4 cells themselves. This is fundamentally different from the ablation approach used here, which removes the S100A4 cell population.

In contrast, studies employing *S100a4-Cre*–driven DTA–mediated ablation represent a directly comparable approach. Such studies have reported systemic phenotypes, including reduced adipose tissue and altered metabolic parameters^26^, indicating that S100A4-expressing cells contribute to multiple aspects of tissue homeostasis. Consistent with these previous reports, *S100a4-Cre;DTA* mice used in our study were significantly smaller than their littermates. Our findings extend these observations by identifying a specific and previously unrecognized role for this cell population in nipple morphogenesis.

Notably, we observed incomplete depletion of S100A4+ cells in the mammary gland and nipple. Interestingly, the same *S100a4-Cre;DTA* mouse model resulted in complete depletion of S100A4+ cells in the superficial layer of the mandibular condyle^47^. This suggests that incomplete depletion of S100A4+ cells in the nipple and mammary gland is due to tissue-specific dynamics, rather than lack of depletion efficiency, indicating a compensatory mechanism that can balance the cell loss.

The transcriptomic data on *S100a4-Cre;DTA* implies the tissue homeostasis perturbation caused directly or indirectly by the constitutive depletion of S100A4+ cells. Most notably, skin exhibits signs of xenobiotic-induced response. While DTA is not a xenobiotic *per se*, it is a foreign exotoxin that triggers cellular responses in the host organism. It is designed to accumulate inside the cell and cause apoptosis without spreading to the surrounding area. For cell-specific depletion studies, attenuated DTA (tox176) is used^48^, where 200 molecules need to accumulate to induce cell death^49^. The time between the onset of *Cre* expression and cell death may vary depending on *Cre* activity, excision efficiency and the metabolic activity of the cell, as DTA blocks protein synthesis^25,50,51^.

The observed tissue response can be also associated with hallmarks of mammary involution, the process which is triggered by the milk stasis. However, the tissues were collected within few hours after parturition, when the effect of involution should be minimal^52^. Rather, we hypothesize that immune cell recruitment, and the upregulation of the lipid skin barrier might be caused in response to the continuous apoptosis of S100A4+ cells and their replacement. The transcriptomic signature also revealed downregulation of genes essential for milk production and luminal epithelial differentiation. This potentially reflects the lack of suckling/milk removal-dependent post-partum alveolar maturation and maintenance of secretory activation^53^.

In summary, this study provides an overview of the heterogeneity of S100A4+ cells and their role in the development of skin derivatives, nipples, and mammary glands. We believe that our work may contribute to advancing research on nipple pathologies, breastfeeding medicine, and lactation.

## Limitations of the study

A limitation of this study is that the timing of DTA-mediated cell depletion cannot be precisely defined in the constitutive mouse model employing *S100a4-Cre* because recombination may occur continuously following the initial expression of *S100a4* (E8.5^18^). This limitation could be overcome by usage of inducible *S100a4-CreERT* instead, which could help determine if the nipple phenotype is established embryonically or during postnatal morphogenesis.

## Materials and Methods

### Animals

All procedures involving animals were performed under the approval of the Ministry of Agriculture of the Czech Republic, supervised by the Expert Committee for Laboratory Animal Welfare at the Faculty of Medicine, Masaryk University (FMMU). Mouse strains *S100a4-Cre* ^24^(stock# 030644), *R26mT/mG* (shortly *“mT/mG”*)^30^ (stock# 037456) and *ROSA26-eGFP-DTA* (shortly “DTA”)^25^ (stock# 006331) were acquired from Jackson Laboratories and maintained on a C57BL/6J background unless specified otherwise. ICR mice were obtained from the animal facility at the FMMU. The experimental animals were obtained by breeding of parental strains (*S100a4-Cre;mT/mG* × *DTA*), the genotypes were determined by genotyping. Mice were kept in individually ventilated cages (breeding pairs or triads), or in open cages (non-breeding mice), with ambient temperature of 22°C, a 12 h:12 h light:dark cycle. They were provided with food and water *ad libitum*.

### Timed-mating experiments

For the study of embryonic development and pregnancy, timed matings were set up for overnight. The morning following a successful mating (determined by the presence of vaginal plug) was considered as day 0.5 of pregnancy/embryogenesis. For the lactation study, adult (16+ weeks old) female *S100a4-Cre;DTA* or *DTA* mice were mated with wild-type males (ICR) to ensure the mating efficiency and good health-condition of the litters. At the time of expected delivery, the mated females were checked every morning. If delivered, the pups were counted and their survival was evaluated, maternal behavior was observed, and tissue was collected in the evening.

### Fostering experiment

On postnatal day 1 (P1), the pups of *S100a4-Cre;DTA* were removed from their nest and placed with a wild-type foster mother, as previously described^54^. The foster mother was monitored to ensure she adopted the pups successfully, and the number of surviving pups was evaluated.

### Determination of estrous cycle stage

Vaginal smears were collected *post mortem* (from freshly culled mice) as a vaginal lavage using 15 µl of PBS, and then spread onto a glass slide. After drying, the samples were processed for Papanicolaou staining^55^. The stained samples were then photographed using a Leica DM5000 microscope equipped with a Leica DFC480 camera. The estrous stage was then determined using the criteria published by Byers and colleagues^56^.

### Whole-mount carmine staining of the mammary glands

Right inguinal mammary glands were collected and spread on microscope slides, fixed with Carnoy’s fixative (100% ethanol:chloroform:glacial acetic acid, 6:3:1) overnight at 4°C, rehydrated, and stained with Carmine Alum [1% (w/v) carmine, 0.5% (w/v) aluminium potassium sulphate] overnight. The following day, the tissue was gradually dehydrated in a series of ethanol baths and cleared in Bio-clear (Bio-Optica). Carmine-stained whole-mounts were photographed using a Leica M165FC microscope equipped with a Leica DFC450C camera. The phenotype analysis was performed using ImageJ. The epithelial outgrowth was quantified as “epithelial outgrowth distally from the lymph node”, i.e., the distance from the middle of the lymph node to the most distal epithelial tip. For early developmental time points, when the epithelium has not yet reached the lymph node, the outgrowth comes out as a negative distance. The fat pad size was measured as the total area of the inguinal mammary gland fat pad.

### Histological staining of formalin-fixed paraffin-embedded (FFPE) mammary and nipple tissue sections

The embryos and newborn pups were euthanized by decapitation. The older mice were euthanized by cervical dislocation at the desired time point. They were weighted, and their tissues were photographed and collected immediately. For the histological analysis, the left inguinal mammary glands were collected and spread on a microscope slide. The tissue was fixed overnight in 10% neutral buffered formalin (NBF) at room temperature (RT). The associated nipples were cut from the skin as a 2 mm diameter patch and fixed overnight in 10% NBF at RT in a tube. After fixation, the tissue was removed from the microscope slides/ tubes, placed in histological cassettes, and washed in running tap water for 1 h. Then, it was processed using a standard paraffin embedding procedure. To prepare histological sections, the paraffin-embedded tissue was cut to a 5 µm thickness, deparaffinized using xylene, and rehydrated in ethanol baths of descending concentrations. Then, the sections were stained with hematoxylin and eosin, Masson’s trichrome or picrosirius red. The stained sections were then photographed using a Zeiss Axioscan 7 microscope, equipped with an Axiocam 705 color R1 camera. To visualize of collagen fiber birefringence in picrosirius red-stained tissue sections, a Zeiss AxioImager.Z2 microscope, equipped with polarized light and an AxioCam 208 camera, was used. The obtained data were exported using ZEN software (Carl Zeiss AG, Germany). Mean Intensity Value of red channel (collagen I) was measured using ZEN software (Carl Zeiss AG, Germany), selecting 3 ROIs per biological replicate in anatomically similar regions. Nipple size was measured in ImageJ as the distance from the nipple-areola ridge to the most distal point. Adipocytes area was measured using QuPath software^57^.

### Collagen fiber analysis

Images of picrosirius red-stained nipple sections were pre-processed using ImageJ. The pre-processing involved background reduction (Otsu Auto-Threshold) and fiber filtering (MorphoLibJ plugin) to mask non-fibrous and bulk structures. The collagen fibers were then analyzed using CT-FIRE v3.0 Beta^58^.

### Immunofluorescent staining of FFPE sections

For the immunofluorescent staining on FFPE sections, the samples were prepared as described above. After deparaffinization and rehydration, the samples were washed with distilled water and incubated for 25 min in 98°C water bath with pH 6 (#S2369, Dako) or pH9 (#S2367, Dako) antigen retrieval buffer. The buffer was selected based on prior optimization for each antibody. After serial washing, the samples were blocked with a blocking buffer [10% foetal bovine serum (FBS; Hyclone/GE Healthcare), and 0.05% Tween-20 (Sigma) in PBS (PBS-T)] for 1 h in a humidified chamber at RT. Then, the samples were incubated with the primary antibody (**Table 1**), diluted in the blocking buffer, overnight in a humidified chamber at 4°C. The next day, the samples were washed with PBS-T and incubated with the secondary antibody in the blocking buffer for 2 h, and with the DAPI solution (1 µg/ml; Merck) for 15 min in a humified chamber at RT, protected from light. Finally, the samples were washed with PBS-T and mounted in Fluoromount-G (Sigma). The tissue sections were photographed using a Zeiss Axio Observer.7 microscope with a LSM 880 laser scanning unit, 405 nm, 488 nm, 561 nm, and 633 nm lasers, a GaAsP detector, and Plan-Apochromat objectives (10x/0.45 AIR, 25x/0.80 MIM, 63x/1.40 with oil immersion; all Zeiss). The obtained data were exported, and the brightness of each channel was linearly enhanced using ZEN software (Carl Zeiss AG, Germany). Cell density, the number of KI67+ cells, and double-stained cells were quantified using Qupath software^57^.

**Table 1.** List of antibodies used in this study.

| **Antibody** | **Type** | **Supplier** | **Cat. Number** | **Dilution** |
|---|---|---|---|---|
| AIF1 | goat polyclonal | Novus Biologicals | NB100-1028 | 1/500 |
| CD31 | Goat polyclonal | Novus Biologicals | AF368 | 1/200 |
| CD45 | rat monoclonal | BD Pharmingen | 550539 | 1/200 |
| E-cadherin | rat monoclonal | Santa Cruz Biotechnology | sc-59778 | 1/50 |
| S100A4 | rabbit polyclonal | Merck Millipore | 07-2274 | 1/200 |
| F4/80 | rat monoclonal | Santa Cruz Biotechnology | sc-52664 | 1/100 |
| GFP | chicken polyclonal | Novus Biologicals | NB100-1614 | 1/300 |
| GFP | chicken polyclonal | Abcam | AB13970 | 1/500 |
| Keratin 5 | rabbit monoclonal | Biolegend | 905504 | 1/500 |
| Keratin 8 | mouse monoclonal | Biolegend | 904804 | 1/500 |
| Keratin 10 | rabbit polyclonal | Biolegend | 905404 | 1/200 |
| KI67 | rabbit monoclonal | Zytomed Systems | RBK027-05 | 1/300 |
| PDGFRα | rabbit polyclonal | Invitrogen | PA5-16571 | 1/100 |
| Vimentin | chicken polyclonal | Invitrogen | PA1-16759 | 1/200 |
| αSMA | mouse monoclonal | eBioscience | 14-9760-82 | 1/300 |
| β-casein | rabbit monoclonal | Santa Cruz Biotechnology | sc-30042 | 1/200 |
| Alexa Fluor 488, 568, or 647-conjugated secondary secondary antibodies | goat secondary<br>donkey secondary | Thermo Fisher Scientific | A11001, A11036, A21449, A11055, A78952 | 1/500 |

### Mammary cryosections

To analyze the *S100a4-Cre;R26mT/mG* native endofluorescence histologically, cryopreserved tissue sections were prepared. First, the thoracic mammary glands were collected, fixed, and washed as previously described. The tissue was then incubated in 15% sucrose at RT with gentle shaking O/N. The next morning it was moved to 30% sucrose and incubated at RT with gentle shaking until the tissue sank. The tissue was then placed into a plastic cryo-mold and embedded in a cryopreservation medium (O.C.T.^TM^ Compound, Tissue-Tek), and snap frozen. The frozen OCT blocks were stored at-80 °C until processed. 7 µm thick tissue cryosections were cut on a Leica CM1850 cryostat, collected on microscopy glass slides, washed in PBS, incubated with DAPI solution (1 µg/ml; Merck), and mounted in Fluoromount-G (Sigma). The tissue sections were photographed using an inverted Zeiss Axio Observer.7 microscope with laser scanning unit LSM 880 using 405 nm, 488 nm, 561 nm lasers, GaAsP detector, and objective Plan-Apochromat 63×/1.40 with oil immersion (Zeiss). The obtained data were exported, and the brightness of each channel was linearly enhanced using ZEN software (Carl Zeiss AG, Germany).

### CUBIC tissue clearing of the mammary gland and nipple

Whole-mount endogenous fluorescence analysis was performed using the optical tissue clearing technique CUBIC^31^. First, the thoracic mammary glands were collected and spread on microscope slides. The nipples were collected as described above. To analyze the mammary glands and nipples of the embryos, the abdominal skin flanks were collected. The tissues were fixed in 10% NBF for 6 h at RT and then washed in PBS. All subsequent steps were performed at RT, with gentle shaking. First, the tissue was immersed in CUBIC1 reagent (25% urea, 25% N,N,N’,N’-tetrakis(2-hydroxypropyl)ethylenediamine, 15% Triton X-100; all Sigma Aldrich) for 3 days. Then, it was transferred to fresh CUBIC 1 for 2 more days. Then, the tissue was washed in PBS and incubated in a DAPI solution (1 µg/ml; Merck) for 2 days. After washing in PBS, the tissues were incubated in CUBIC2 reagent (50% sucrose, 25% urea, 10% 2,2°,2°,-nitrilotriethanol, 0.1% Triton X-100; all Sigma Aldrich) for 2 days. The tissue was then mounted between two coverslips in CUBIC2 reagent. Whole-mounts were photographed using an inverted microscope Zeiss Axio Observer.7 with LSM 880 laser scanning unit using 405 nm, 488 nm and 561 nm lasers, a GaAsP detector, and Plan-Apochromat objectives 10x/0.45 AIR, Plan-Apochromat 25x / 0.80 MIM with oil immersion (all Zeiss). The obtained data were exported and the brightness of each channel was linearly enhanced using ZEN software (Carl Zeiss AG, Germany).

### Oxytocin stimulation assay

For the *ex vivo* oxytocin stimulation assay, the thoracic mammary gland was collected and immediately cut into pieces approximately 5 × 5 mm with scalpels. The pieces were transferred to pre-warmed DMEM/F12 with 10% FBS and incubated at 37 °C until imaged. For imaging and stimulation, a tissue sample was placed in a 35-mm imaging dish (Ibidi) containing 200 µl of PBS. Imaging was performed using an inverted Zeiss Axio Observer.7 microscope with a LSM 880 laser scanning unit, a 561 nm laser, a GaAsP detector, and a Plan-Apochromat 25×/0.80 MIM oil immersion objective (all Zeiss). For time-lapse imaging, one “z” plane was imaged, at a time-frame cycle of 1.88 s. After a brief imaging period without stimulation (approximately 10 time frames), 200 µl of 150 nM oxytocin (Sigma O3251) was pipetted onto the tissue, creating a final oxytocin concentration pf 75 nM. Time-lapse imaging continued without interruption, and the tissue was imaged for approximately 300 time frames. Responsiveness to oxytocin stimulation was evaluated based on the presence of alveolar contractions.

### RNA-seq sample collection

Patches of skin containing thoracic and inguinal nipples, approximately 2 mm in diameter, were dissected from L1 lactating dams and transferred to QIAzol Lysis Reagent (Qiagen). All samples were stored at-80 °C until the complete experimental sample set was collected.

### RNA isolation

RNA was isolated from frozen tissue using 1.4 mm Ceramic Bead Media (Revity), QIAzol Lysis Reagent (Qiagen), and a NucleoSpin RNA kit (Macherey-Nagel). First, the tissue was disrupted in a homogenizer (Bead Ruptor 4, Omni International) with 800 μl of Qiazol and ceramic beads. Then, the aqueous phase containing RNA was separated using 80 μl of BCP (Molecular Research Center). The aqueous phase was pipetted out and mixed with 70% ethanol in a 1:1 ratio. Then, it was applied to the NucleoSpin RNA column. The rest of the isolation was performed according to the NucleoSpin RNA kit manufacturer’s protocol. The RNA concentration was determined using a Nanodrop 2000c (Thermo Scientific), and the RNA integrity was checked using a Fragment Analyzer and an RNA Kit 15 nt (Agilent Technologies).

### QuantSeq 3’mRNA-seq library prep

500 ng of total RNA was used as the input for the library preparation, which was performed using the QuantSeq 3′ mRNA-Seq V2 Library Prep Kit FWD with UDI 12 nt (Lexogen), in combination with UMI Second Strand Synthesis Module for QuantSeq FWD (Lexogen). The library’s quantity and size distribution were checked using the QuantiFluor dsDNA System (Promega) and the High Sensitivity NGS Fragment Analysis Kit (Agilent Technologies). The final library pool was sequenced using an AVITI 2×75 Sequencing Kit Cloudbreak FS High Output (Element Biosciences) on an AVITI System (Element Biosciences), resulting in an average of 11 million reads per sample.

### RNA-seq analysis

The *Mus musculus* reference genome, version GRCm39, was downloaded from the Genome Reference Consortium (https://www.ncbi.nlm.nih.gov/grc/mouse). Read quality was assessed using FastQC version 0.11.9 (https://www.bioinformatics.babraham.ac.uk/projects/fastqc) and MultiQC version 1.18^59^. Low-quality base pairs, polyadenine tails and adapter contamination were removed using Cutadapt version 4.4^60^. Clean reads were then aligned to the genome sequence with HISAT2, version 2.2.1^61^, and SAMtools, version 1.13^62^. The aligned reads were then deduplicated using UMI-tools version, 1.1.6 ^63^. The alignment was then inspected visually in the IGV browser, version 2.9^64^. Gene counts were generated using the GenomicAlignments package ^65^ in R/Bioconductor (https://R-project.org)^66^. Differentially expressed genes (DEG) were detected using DESeq2^67^. Gene ontology term enrichment analysis (https://geneontology.org) was performed using the gseGO and enrichGO functions in clusterProfiler^68^.

### Statistical analysis

The sample size was not determined *a priori*, and the investigators were not blinded to the experimental conditions. Calculations were performed using Prism 6 (GraphPad). The Mann-Whitney test and one-way ANOVA was used to determine statistical significance. All data are presented as the mean ± standard deviation (SD). ns P > 0.05, *P < 0.05, **P < 0.01, ***P < 0.001, and ****P < 0.0001. The number of independent biological replicates is indicated as n, and the number of individual samples is indicated as N.

## Data Availability Statement

RNA-seq data are available at ArrayExpress database (https://www.ebi.ac.uk/arrayexpress/) under the accession number: E-MTAB-16579.

## Author Contributions

D.J.B. managed mouse colony, performed most experiments, analyzed data, prepared figures, and wrote the manuscript. E.G. performed experiments and analyzed mammary gland phenotypes. V.Z. analyzed bulk RNA sequencing data. Z.S.K. conceptualized the study, managed mouse colony, and wrote the manuscript. D.J.B. and Z.S.K. acquired the funding for the study. All authors reviewed and edited the final manuscript.

## Acknowledgements

We thank Katarina Mareckova for technical support with tissue processing for histology. We thank David Vondrasek for assistance with collagen fiber analysis, including pre-processing design and guidance on analytical approaches. We thank Matea Brezak for her help with mouse husbandry. We acknowledge the core facility (CF) CELLIM of CEITEC supported by the Czech-BioImaging large RI project (LM2023050), funded by MEYS CR, and the CF Genomics and the CF Bioinformatics supported by the NCMG research infrastructure (LM2023067 funded by MEYS CR) for their support with obtaining scientific data presented in this paper.

## Abbreviations

DTA: diphtheria toxin fragment A
FOV: field of view
GFP: green fluorescent protein
KRT5: keratin 5
KRT8: keratin 8
mT: membrane tomato
PDGFRα: platelet-derived growth factor receptor α
αSMA: α smooth muscle actin
VIM: vimentin

