## Supplementary Material for "S100A4+ stromal cells are required for functional nipple development and successful nursing"

**Supplementary Figures**

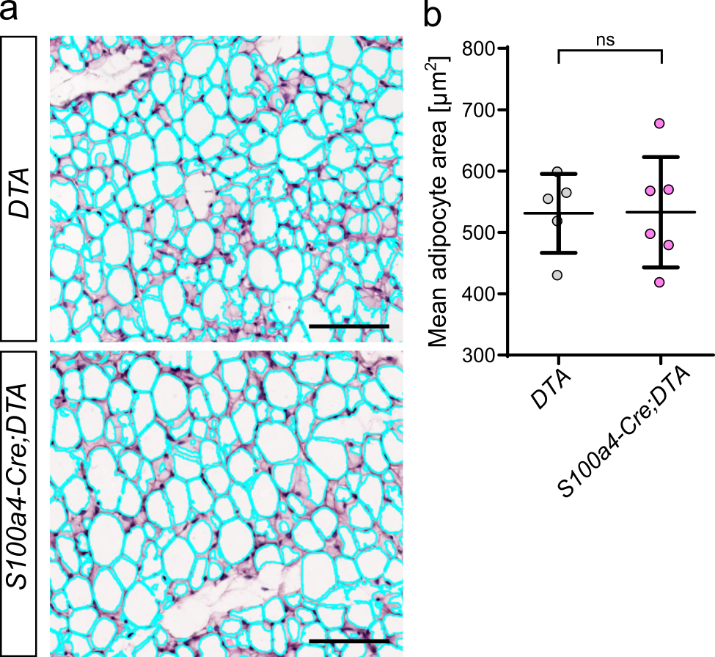

**Figure S1. Adipocyte size is mantained in *S100a4-Cre;DTA* mice.** (a) Representative images of adipocyte area analysis performed on H&E-stained FFPE sections of the distal fat pad from 7-week-old *DTA* and *S100a4-Cre;DTA* mice. Cyan outlines delineate individual adipocytes. Scale bar = 100 µm. (b) Quantification of mean adipocyte area. The plots show the mean ± SD, ns p > 0.05, (Mann-Whitney test), n = 5 *DTA*/ 6 *S100a4-Cre;DTA*, each dot represents the mean value for one biological replicate.

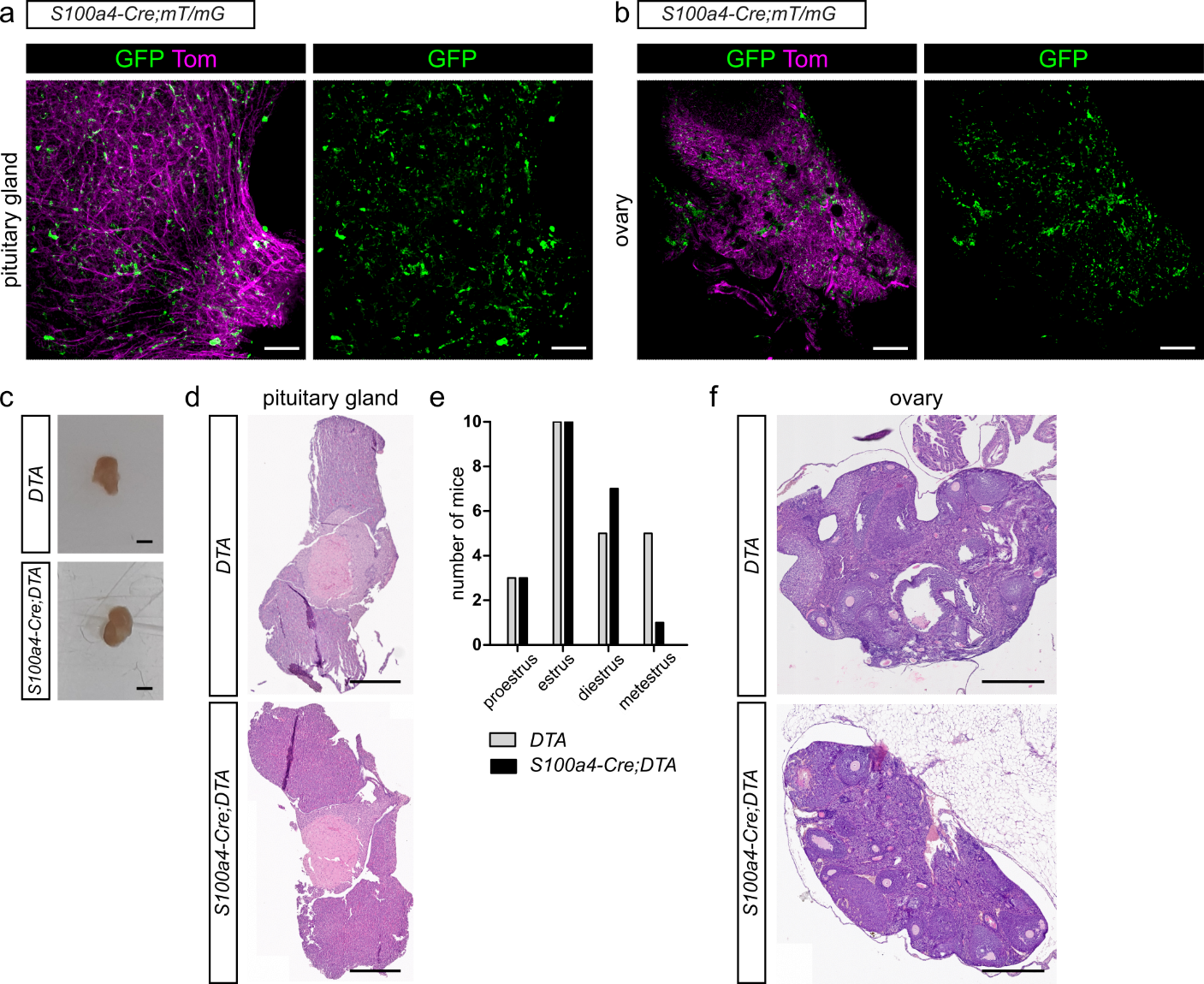

**Figure S2. Hormone-producing organs, ovary and pituitary gland, have a physiological structure in *S100a4-Cre;DTA* mice.** (a) Representative images of S100A4-lineage cells in cleared whole-mount *S100a4-Cre;mT/mG* pituitary gland. Scale bar = 100 µm. (b) Representative images of S100A4-lineage cells in cleared whole-mount *S100a4-Cre;mT/mG* ovary. Scale bar = 100 µm. (c) Macroscopic images of unprocessed *DTA* and *S100a4-Cre;DTA* pituitary glands after dissection. Scale bar = 1 mm. (d) H&E-stained FFPE tissue sections of *DTA* and *S100a4-Cre;DTA* pituitary glands. Scale bar = 500 µm. (e) Graphical presentation of the estrus cycle stage distribution among adult *DTA* and *S100a4-Cre;DTA* mice. N = 23 *DTA*/ 21 *S100a4-Cre;DTA*. (f) H&E-stained FFPE tissue sections of a *DTA* and a *S100a4-Cre;DTA* ovary. Scale bar = 100 µm.

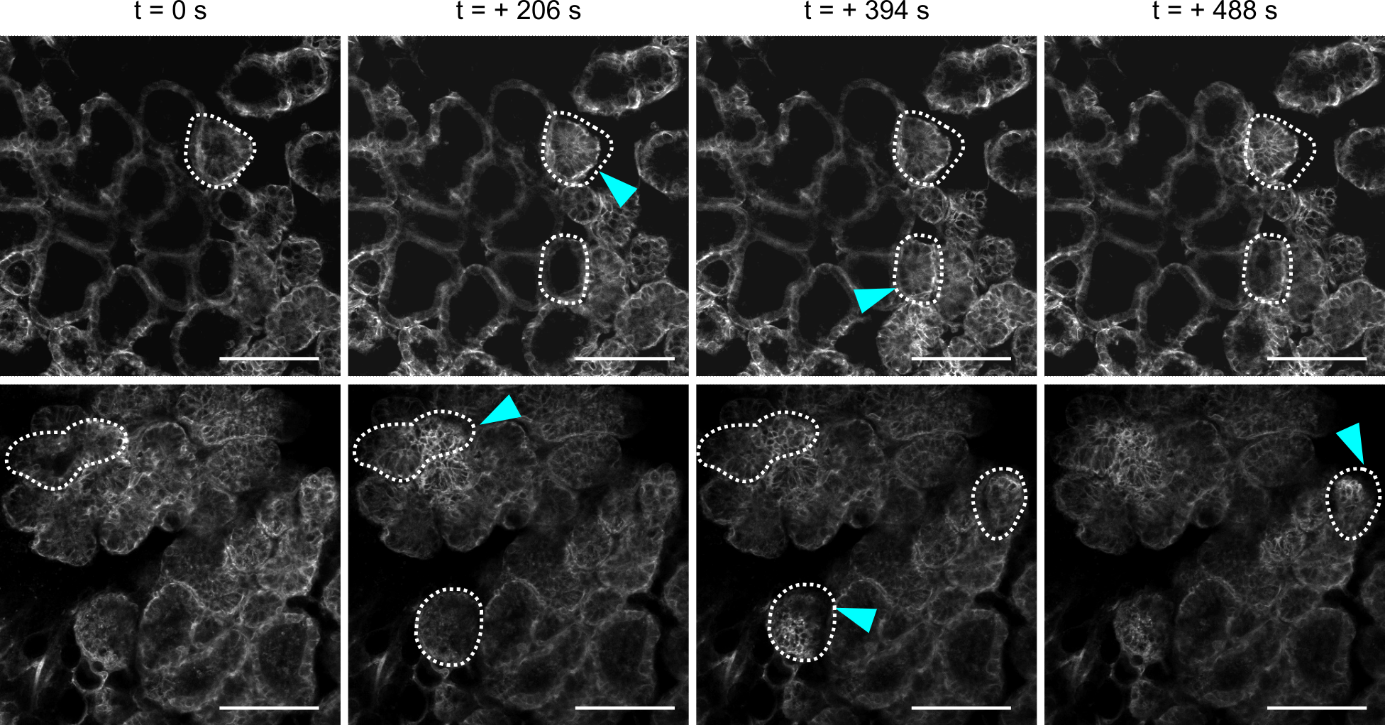

**Figure S3. Response to oxytocin stimulation is retained in lactating mammary gland from *S100a4-Cre;DTA* mice.** Snapshots from time-lapse imaging of ex-vivo oxytocin stimulation assay on lactating mammary glands from *DTA* and *S100a4-Cre;DTA* mice. Dotted lines encircle the most visible contracting alveoli and arrowheads pinpoint contraction events. Scale bar = 100 µm. See Movies S1 and S2 for the full sequence.

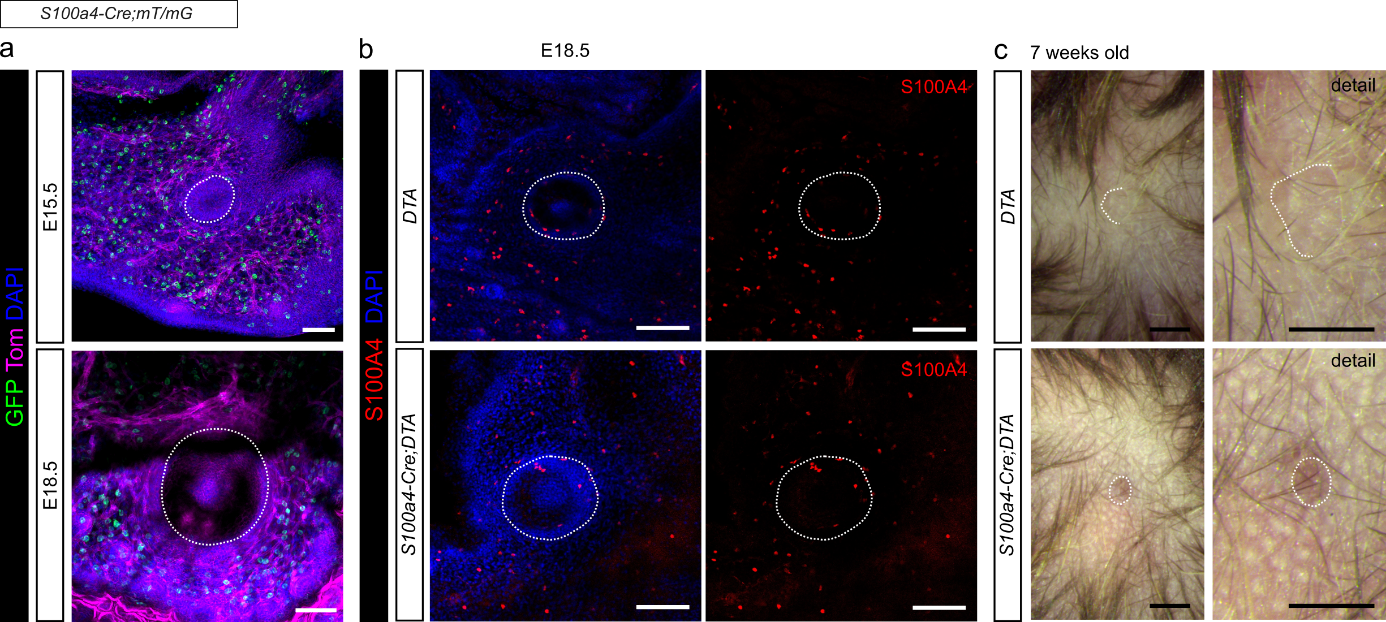
**Figure S4. Embryonic and pubertal nipple phenotypes.** (a) Representative images of cleared whole-mount *S100a4-Cre;mT/mG* nipple tissue at embryonic developmental time-points: E15.5 and E18.5. Scale bar = 100 µm. (b) Immunofluorescent labeling for S100A4 on embryonic *DTA* and *S100a4-Cre;DTA* whole-mount skin (E18.5). Scale bar = 100 µm. (c) Representative in situ photographs of nipples from DTA and *S100a4-Cre;DTA* pubertal (7-weeks old) mice. Scale bar = 1 mm. Dotted lines indicate nipple border.

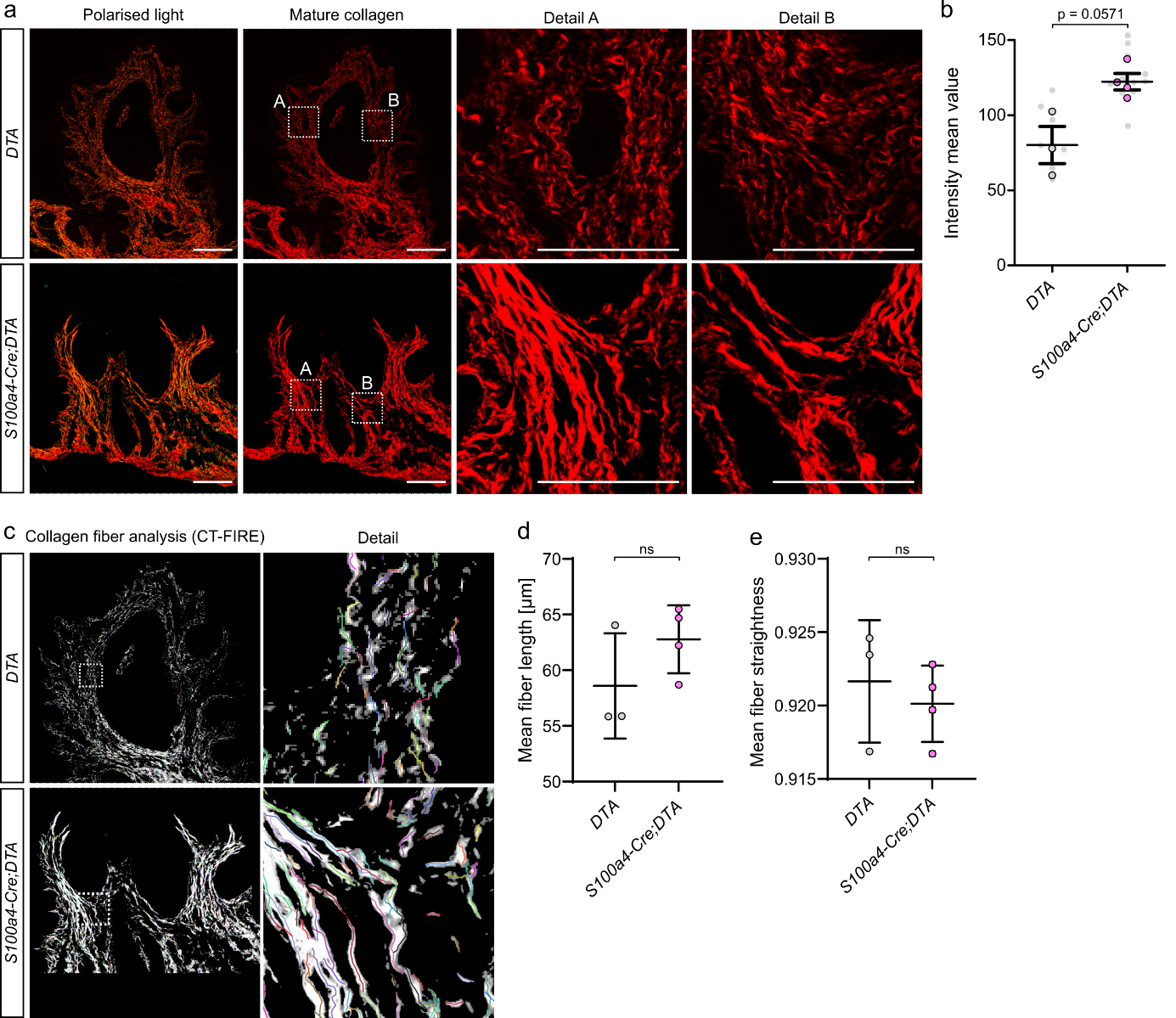

**Figure S5.** **Collagen fibers are densely packed in *S100a4-Cre;DTA* nipples.** (a) Representative pictures of histological sections of *DTA* and *S100a4-Cre;DTA* nipples stained for collagen by picrosirius red and imaged in polarised light to detect dense collagen fibers. The red channel (collagen I) is shown alongside details of selected regions A and B. Scale bars = 200 µm and 100 µm (in detail pictures). (b) Quantification of Intensity mean value for the red channel (dense collagen). (c) Representative images of CT-FIRE collagen fiber analysis on *DTA* and *S100a4-Cre;DTA* nipples. (d, e) Quantification of collagen fiber length (d) and straightness (e) from CT-FIRE analysis. The plots in (b, d, e) show mean ± SD, ns p > 0.05 (Mann-Whitney test), n = 3 *DTA* / 4 *S100a4-Cre;DTA*, each dot represents a mean value per biological replicate.

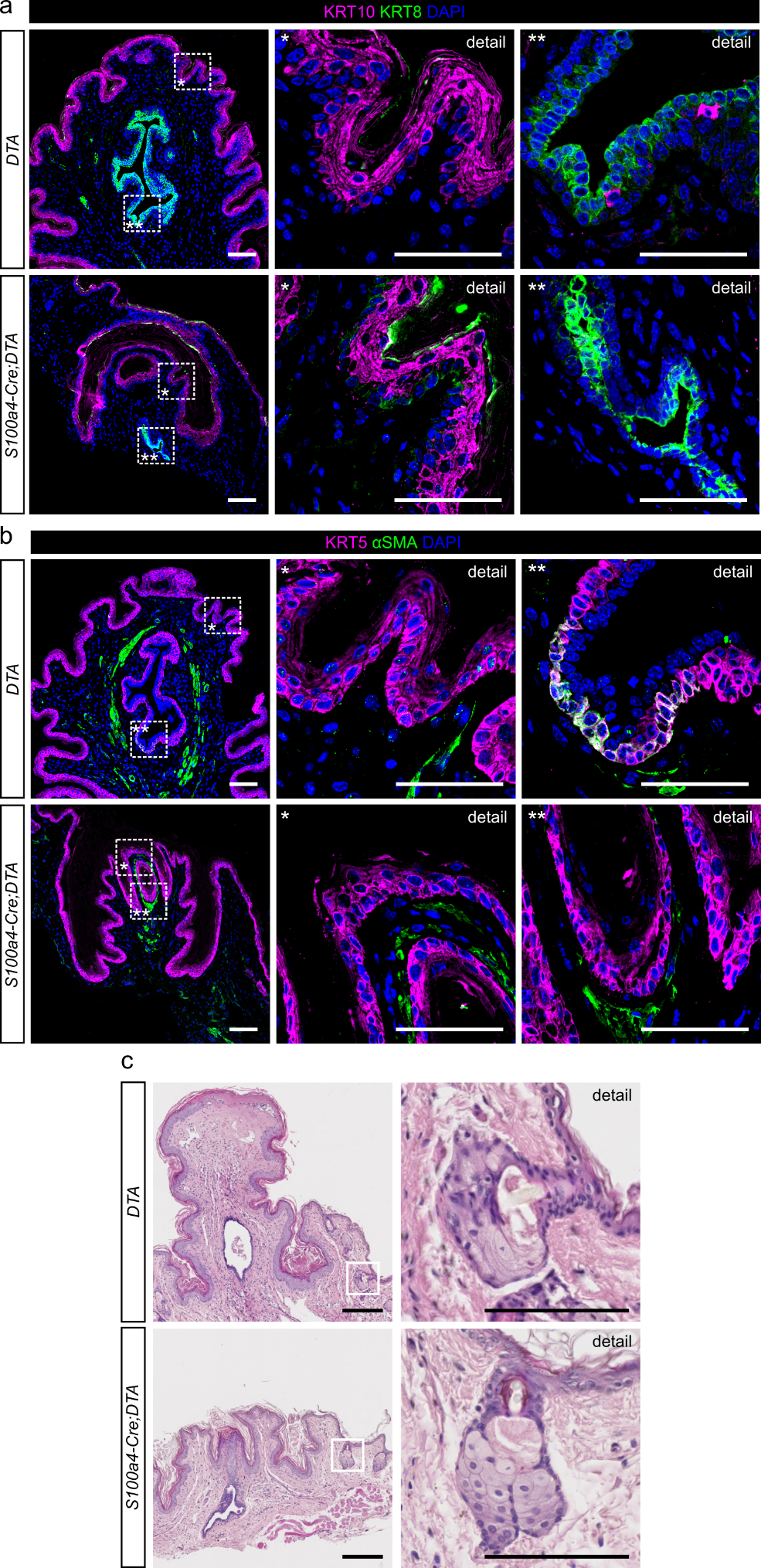

**Figure S6. Nipple tissue architecture is unaltered in *S100a4-Cre;DTA* mice.** (a) Immunofluorescent labeling for KRT8 and KRT10 on FFPE sections of *DTA* and *S100a4-Cre;DTA* nipples. (b) Immunofluorescent labeling for KRT5 and αSMA on FFPE sections of *DTA* and *S100a4-Cre;DTA* nipples. Bands of smooth muscle extend along the lactiferous duct in both. Scale bars (a, b) = 100 µm. (c) H&E-stained FFPE tissue sections of the *DTA* and *S100a4-Cre;DTA* nipples are focused on sebaceous glands. The white boxes in the lower magnification pictures mark tissue area shown in the higher-magnification pictures. Scale bars = 200 µm and 100 µm (detail).

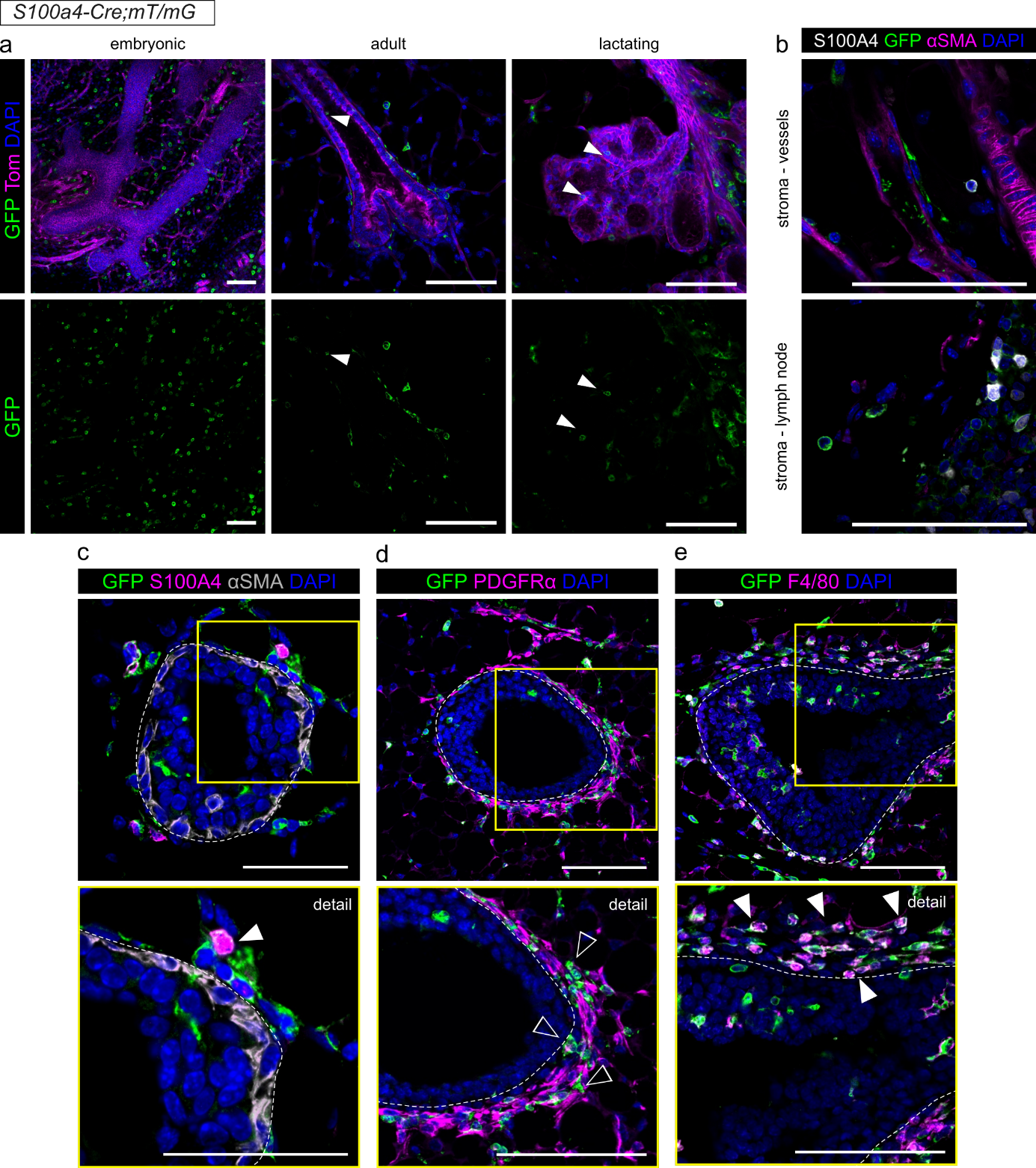

**Figure S7. S100A4+ lineage cells are a heterogeneous stromal cell population in the mammary gland.** (a) Representative images of *S100a4-Cre;mT/mG* cleared whole-mount mammary glands at various developmental time points: embryonic (E18.5), adult virgin (10 weeks of age), and lactation (L1). Arrowheads pinpoint GFP+ cells localized within mammary epithelium. (b) Immunofluorescent labeling of S100A4, αSMA and GFP on *S100a4-Cre;mT/mG* FFPE mammary gland sections. Representative images depict different stromal locations (vessel and lymph node), highlighting the heterogeneity of S100A4+ cell lineage. (c-e) Immunofluorescent co-staining of GFP and S100A4 (c), PDGFRα (d) and F4/80 (e) on FFPE sections of *S100a4-Cre;mT/mG* mammary gland tissue. Dotted line encircles the epithelium. White arrowheads highlight the double-stained cells, empty arrowheads point to single-stained GFP+ cells, yellow boxes mark the area depicted in the detailed picture of higher magnification. Scale bar = 50 µm (yellow boxes with detail) and 100 µm.

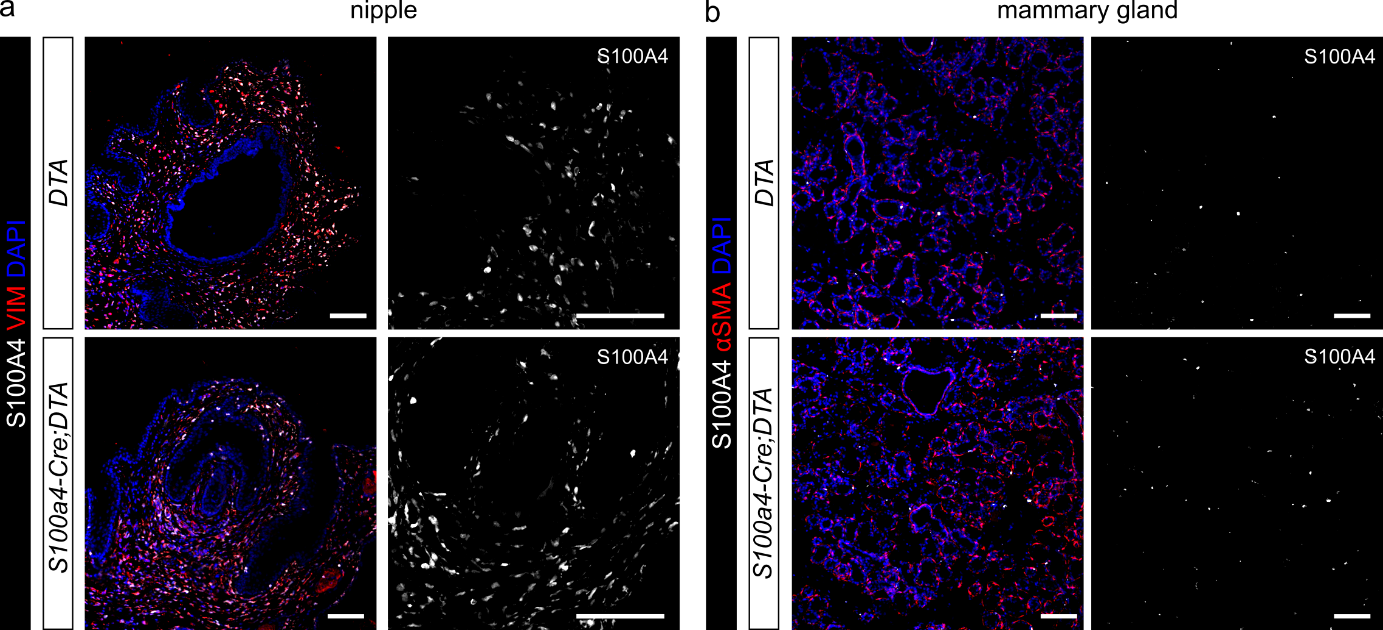

**Figure S8. S100A4+ cells are found in *S100a4-Cre;DTA* nipple and mammary tissues.** (a) Immunofluorescent labeling for S100A4 and vimentin on FFPE sections of *DTA* and *S100a4-Cre;DTA* L1 nipples. Scale bar = 100 µm. (b) Immunofluorescent labeling for S100A4 and αSMA on FFPE sections of *DTA* and *S100a4-Cre;DTA* L1 mammary gland. Scale bar = 100 µm.

**Supplementary Movies**

**Movie S1.** ***DTA* lactating mammary tissue contracts in response to oxytocin stimulation.** Time-lapse video of ex-vivo oxytocin stimulation assay on a lactating mammary tissue from *DTA* mouse.

**Movie S2. Lactating mammary tissue from *S100a4-Cre;DTA* mice shows an unaltered response to oxytocin stimulation.** Time-lapse video of ex-vivo oxytocin stimulation assay on a lactating mammary tissue from *S100a4-Cre;DTA* mouse.
